# Proteobacteria encode diverse flg22 peptides that elicit varying immune responses in *Arabidopsis thaliana*

**DOI:** 10.1101/2020.11.09.374983

**Authors:** Janis H.T. Cheng, Melissa Bredow, Jacqueline Monaghan, George C. diCenzo

**Author notes:** These authors contributed equally to this work (Janis H.T. Cheng, Melissa Bredow).

## Abstract

Bacterial flagellin protein is a potent microbe-associated molecular pattern. Immune responses are triggered by a 22 amino acid epitope derived from flagellin, known as flg22, upon detection by the pattern recognition receptor FLAGELLIN-SENSING2 (FLS2) in multiple plant species. However, increasing evidence suggests that flg22 epitopes of several bacterial species are not universally immunogenic to plants. We investigated whether flg22 immunogenicity systematically differs between classes of the phylum Proteobacteria, using a dataset of 2,470 flg22 sequences. To predict which species encode highly immunogenic flg22 epitopes, we queried a custom motif (^11^[ST]xx[DN][DN]xAGxxI^21^) in the flg22 sequences, followed by sequence conservation analysis and protein structural modelling. These data led us to hypothesize that most flg22 epitopes of the γ- and β-Proteobacteria are highly immunogenic, whereas most flg22 epitopes of the α-, δ-, and ε-Proteobacteria are weakly to moderately immunogenic. To test this hypothesis, we generated synthetic peptides representative of the flg22 epitopes of each proteobacterial class, and we monitored their ability to elicit an immune response in *Arabidopsis thaliana*. Flg22 peptides of the γ- and β-Proteobacteria triggered strong oxidative bursts, whereas peptides from the ε-, δ-, and α-Proteobacteria triggered moderate, weak, or no response, respectively. These data suggest flg22 immunogenicity is not highly conserved across the phylum Proteobacteria. We postulate that sequence divergence of each taxonomic class was present prior to the evolution of FLS2, and that the ligand specificity of *A. thaliana* FLS2 was driven by the flg22 epitopes of the γ- and β-proteobacteria, a monophyletic group containing many common phytopathogens.

## MAIN TEXT

Plants live in microbe-rich environments and have adapted complex relationships with their microbiota. Mutualistic plant-microbe interactions have placed strong selective pressures on plants to accommodate microbial communities that enhance the uptake of nutrients and contribute to plant fitness. At the same time, pathogenic plant-microbe interactions have placed strong selective pressures on plants to evolve a robust immune system capable of sensing and responding to danger. Key to this sense-and-response system are integral membrane receptor complexes that bind microbial ligands (‘molecular patterns’) with high affinity and trigger phosphorylation-mediated signal transduction cascades (Couto and Zipfel 2016). Although often referred to as ‘pathogen’ associated molecular patterns (PAMPs), some scholars prefer the term ‘microbe’ associated molecular patterns (MAMPs), as these features are shared by entire classes of microbes including pathogens, commensals, and mutualists (Ausubel 2005). How plants differentiate friend from foe, despite all groups of microbes containing MAMPs, remains unclear but is likely to involve complex, overlapping mechanisms (Plett and Martin 2018). This discrimination may depend on concurrent or subsequent release of plant-dervied signals such as cellular debris or phytocytokines that bind additional receptors and amplify immune signaling (Gust et al. 2017). Pathogens and non-pathogens may also directly suppress immune signalling to gain access to plant hosts, and/or encode divergent MAMPs that evade detection by plant receptors (Plett and Martin 2018; Gourion et al. 2015).

One of the most well-known MAMPs is the bacterial flagellin protein (Felix et al. 1999). In several plant species, orthologs of the FLAGELLIN-SENSING 2 (FLS2) receptor detect flagellin via a 22 amino acid epitope known as flg22 (Felix et al. 1999; Gómez-Gómez and Boller 2000; Nicaise et al. 2009). Flg22 is thought to be a broadly conserved and potent elicitor of plant immunity, based predominantly on studies using flagellin peptides of the genus *Pseudomonas* (class γ-Proteobacteria) (Felix et al. 1999). However, there is increasing evidence that the immunogenicity of flg22 epitopes differs between taxonomically diverse bacteria and is not necessarily correlated with bacterial lifestyle. For example, flg22 epitopes encoded by the the plant pathogen *Xanthomonas campestris* (γ-Proteobacteria) and the plant commensal *Paraburkholderia phytofirmans* (β-Proteobacteria) elicit an immune response in *Arabidopsis thaliana*, whereas flg22 epitopes from the plant mutualist *Sinorhizobium meliloti* (α-Proteobacteria), and the plant pathogens *Agrobacterium tumefaciens* (α-Proteobacteria) and *Ralstonia solanacearum* (β-Proteobacteria) do not induce an immune response in *A. thaliana* and/or tomato (Felix et al. 1999; Bauer et al. 2001; Pfund et al. 2004; Sun et al. 2006; Trdá et al. 2014). These observations raise the possibility that the immunogenic properties of flg22 are not broadly conserved and that bacterial lifestyle is not a major driver of flg22 evolution.

To investigate the evolution of flg22 and its immunogenic properties, we performed an *in silico* survey of proteobacterial flg22 epitopes to predict which are likely to be highly immunogenic to *A. thaliana*. To accomplish this, we collected the amino acid sequences of 2,470 flagellin proteins from a representative set of 1,414 Proteobacteria (366 α-Proteobacteria, 234 β-Proteobacteria, 674 γ-Proteobacteria, 74 δ-Proteobacteria, and 66 ε-Proteobacteria), and extracted flg22 sequences based on multiple sequence alignments and the positions of known flg22 epitopes (**File S1 and Dataset S1**). This process yielded 2,404 peptides, encompassing 1,059 distinct peptide sequences. As an initial prediction of which flg22 epitopes might be highly immunogenic to *A. thaliana*, we queried the sequences for the motif ^11^[ST]xx[DN][DN]xAGxxI^21^, which includes amino acids important for activation of the FLS2 - BRASSINOSTEROID INSENSITIVE1-ASSOCIATED KINASE1 (BAK1) immune signaling complex (Felix et al. 1999; Sun et al. 2013; Wei et al. 2020). We allowed for Ser or Thr at residue 11 as these amino acids have similar properties, while the flexibility of Asp or Asn at residues 14 and 15 was based on their functional interchangeability at residue 14 in the flg22 epitope of *X. campestris* (Sun et al. 2006). The ^11^[ST]xx[DN][DN]xAGxxI^21^ motif was found in the flg22 epitopes of > 90% γ- and β-Proteobacterial species that encode flagellin. In contrast, this motif was identified in the flg22 epitopes of only 60%, 15%, and 2% of flagellin-encoding δ-, α-, and ε-Proteobacteria, respectively (**Table S1, Figures 1, S1, and S2**). This result suggested that highly immunogenic flg22 epitopes are likely to be unevenly distributed across the Proteobacteria.

**Figure 1.**
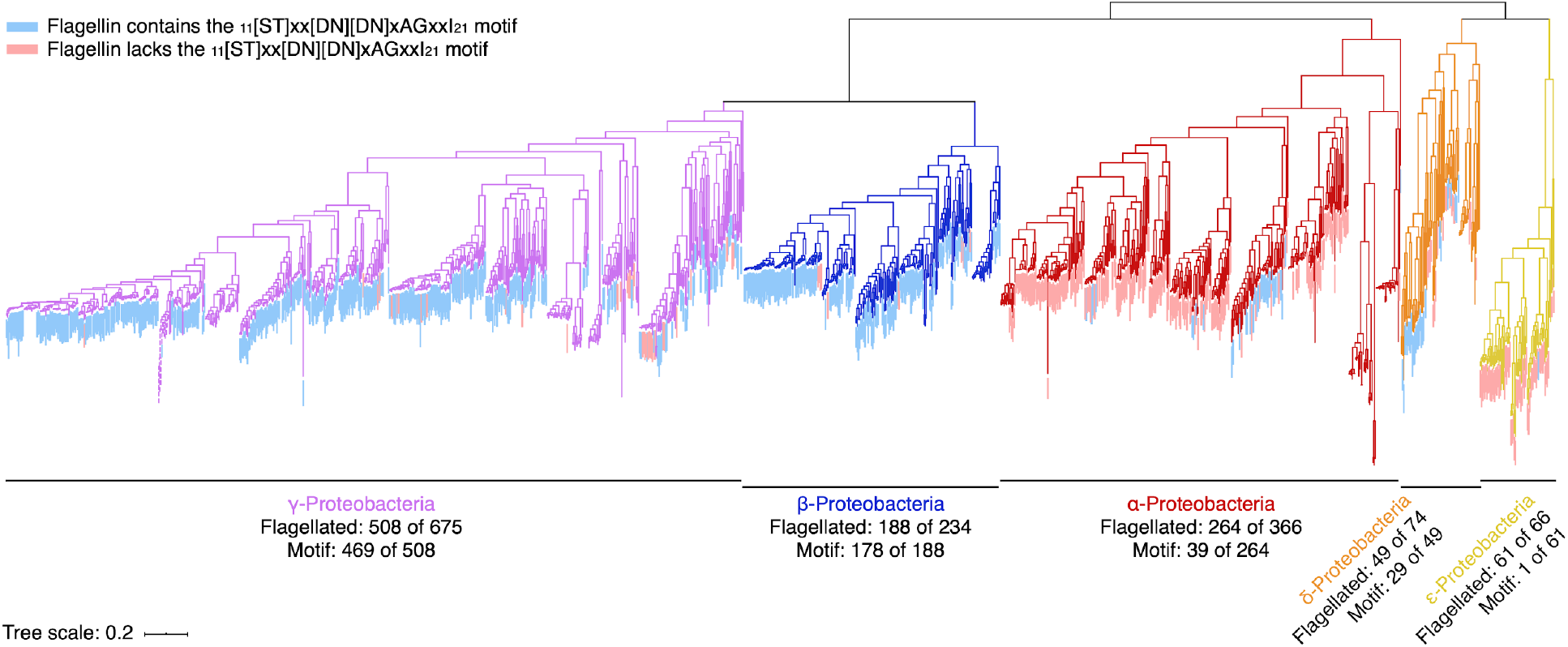
Distribution of flagellin within the Proteobacteria. A maximum likelihood phylogeny of 1,414 Proteobacteria is shown, which was prepared from a concatenated alignment of 31 highly conserved proteins. Branches on the tree are colour coded based on taxonomic class. Taxa encoding a flagellin containing the ^11^[ST]xx[DN][DN]xAGxxI^21^ motif within flg22 are shown in blue; taxa only encoding flagellin that lack this motif are shown in pink, while taxa lacking flagellin are not coloured. For each taxonomic class, the number of species containing flagellin is shown, as is the number of species containing the above-mentioned motif. The scale bar represents the mean number of amino acid substitutions per site. The lengths of branches of a clade of γ-proteobacterial endosymbionts (dashed lines) were reduced for presentation purposes. A larger version of this figure, with legible taxa names, is provided as Figure S1.

This above suggestion was further supported by analysis of flg22 amino acid sequence conservation both within and across taxonomic classes (**Figure 2**), and by *in silico* modelling of the interactions between representative flg22 peptides of each proteobacterial class (herein defined as consisting of the most commonly occurring amino acid at each position [**Figure 2**], and termed flg22-γ, flg22-β, flg22-α, flg22-δ, flg22-ε), in complex with FLS2 and/or BAK1 (**Figures 3 and S3**). Analysis of protein interactions was carried out using the crystal structure of the flg22-bound FLS2/BAK1 complex (Sun et al. 2013) and visualized using PyMOL Molecular Graphics System Version 2.0 Schrödinger, LLC. MutaBind2 (Zhang et al. 2020) was used to predict changes in flg22 peptide binding affinity to FLS2 or binding affinity of flg22-bound FLS2 to BAK1 to simulate the two-step complex formation (Chinchilla et al. 2007; Sun et al. 2013) (**Table S2**). Flg22-γ and flg22-β differed at four and three positions, respectively, from the canonical flg22 sequence of *Pseudomonas aeruginosa* (flg22-Pae) known to induce a strong oxidative burst in several plants including *A. thaliana*, tomato, and grapevine (Felix et al. 1999; Trdá et al. 2014). Nevertheless, flg22-γ and flg22-β were predicted to bind the FLS2/BAK1 receptor complex with high affinity (**Table S2**). In contrast, nine of the flg22-α residues differed from flg22-Pae, and a strong decrease in binding to FLS2 (ΔΔG_bind_=2.37 kcal/mol) and BAK1 (ΔΔG_bind_=1.79 kcal/mol) was predicted. Flg22-δ differed from flg22-Pae at seven residues, although no impact on binding to FLS2 or BAK1 was predicted. However, formation of the flg22-FLS2-BAK1 complex could be negatively impacted by the Arg at position 22, which is positively charged and larger than the Ser or Ala in flg22-γ or flg22-β; as residue 22 is missing from the crystal structure (Sun et al. 2013), we were unable to test this hypothesis *in silico*. Flg22-ε differed from flg22-Pae at eight residues. Binding of flg22-ε to FLS2 was predicted to be moderately decreased (ΔΔG_bind_=0.81 kcal/mol) compared to flg22-Pae, with protein structural modelling suggesting this may be partially due to slight steric hindrance caused by the Ser-17 in flg22-ε (**Figure 3C**).

**Figure 2.**
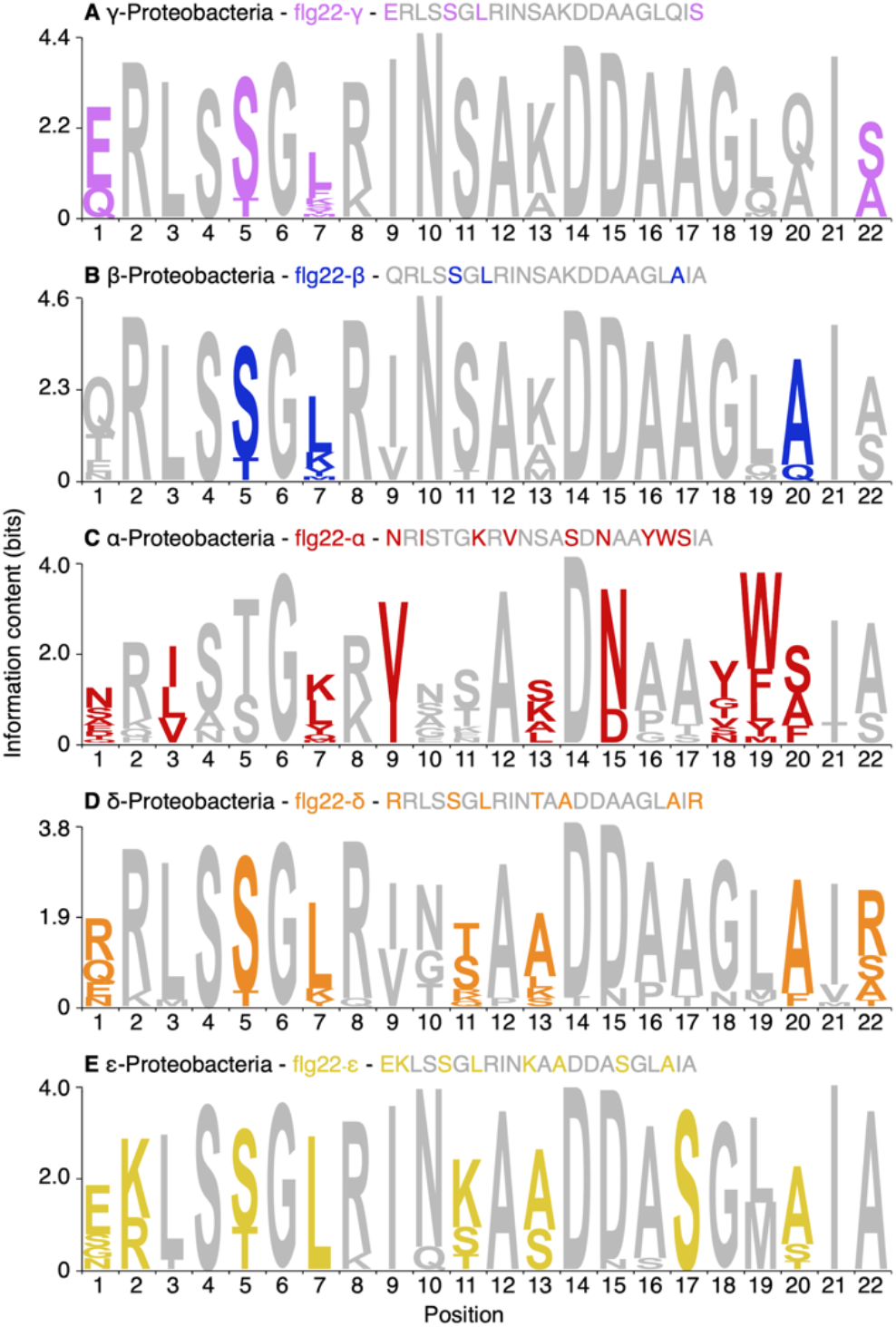
Sequence conservation of the flg22 epitopes of the Proteobacteria. Amino acid logos of the flg22 epitopes of five proteobacterial classes are shown. Logos were generated using Skylign (Wheeler et al. 2014) based on all of the flg22 epitopes identified in each taxonomic class. For each logo, colours indicate the positions that differ (defined as having a different most common amino acid at that position) from the flg22 sequence of *P. aeruginosa*. The representative sequence (defined as consisting of the most commonly occurring amino acid at each position) of each taxonomic class is given above each logo.

**Figure 3.**
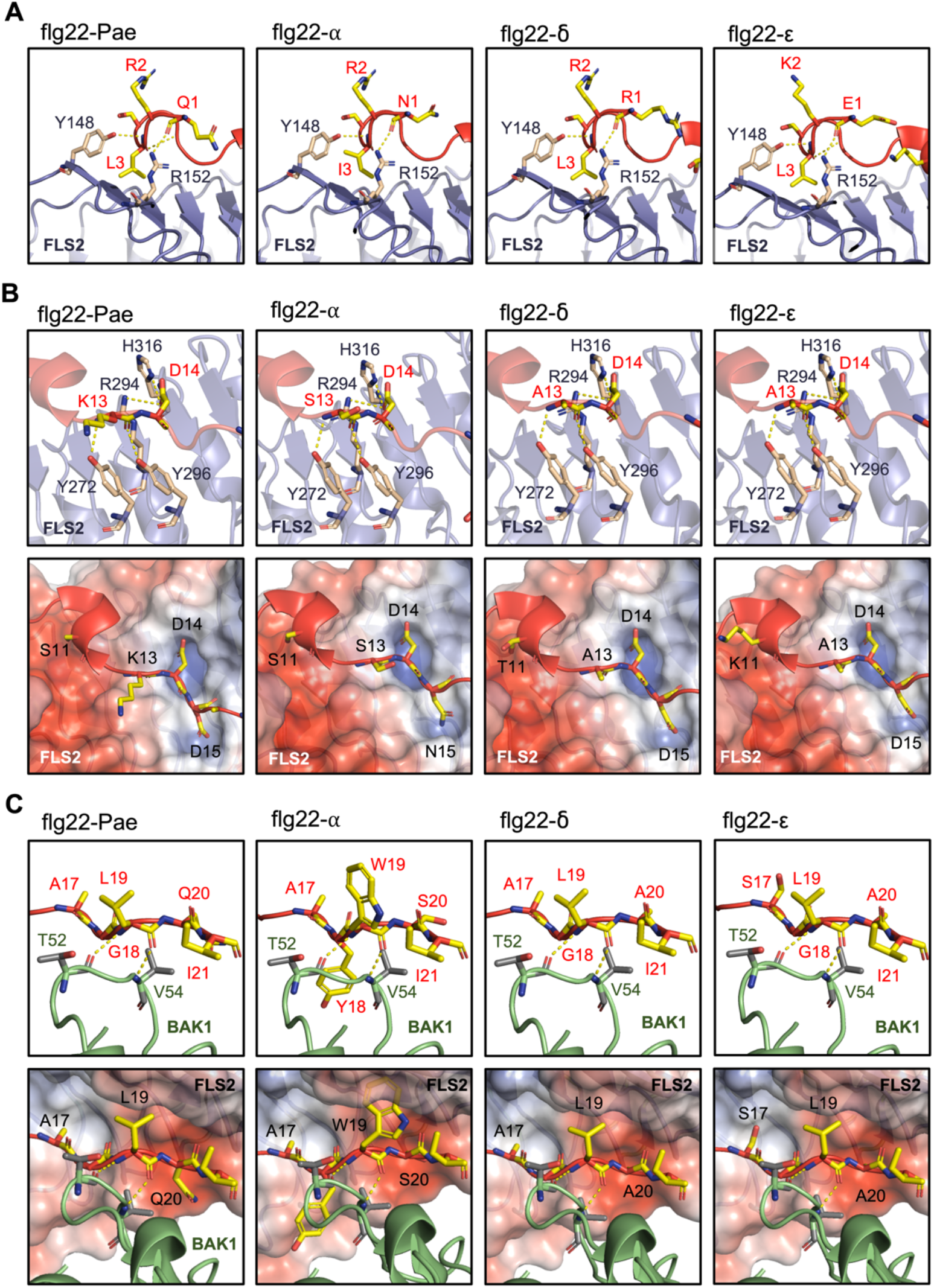
Structural modelling of the FLS2-flg22-BAK1 complex using peptide sequences derived from α-, δ- and ε-Proteobacteria. (**A**) Association between the amino-terminal portion of flg22 (positions 1-9) (red) and FLS2 (purple). The flg22 epitope encoded by *P. aeruginosa* (Pae) and FLS2 is stabilized by H-bonds (dotted yellow lines) between the backbone of flg22 at Gln-1 and Leu-3 and the side chains of Tyr-148 and Arg-152 of FLS2, respectively (Sun et al. 2013). H-bonding is predicted at these positions for all modelled peptides. (**B**) Interaction between the central portion of flg22 (positions 10-15) and FLS2 is stabilized by H-bonds between Tyr-272, Arg-294, Tyr-296, and His-316 of FLS2 and Leu-13 and Asp-14 of flg22 (Sun et al. 2013), which is predicted for all flg22 peptides. Asp-14 and Asp-15 of flg22-Pae interact with two positively charged pockets on the surface of FLS2 (blue) (Sun et al. 2013). Substitution of Asp-15 with Asn-15 in flg22-α is not expected to impact interaction with FLS2 (Sun et al. 2006). (**C**) The carboxyl-terminal portion of flg22 (positions 16-22) interacts with FLS2 and BAK1 (green). Leu-19 of flg22 forms H-bonds with Thr-52 and Val-54 of BAK1 (Sun et al. 2013) and interacts with a hydrophobic pocket (red) on the surface of FLS2 (Sun et al. 2013). Tyr and Trp residues at positions 18 and 19 of flg22-α, respectively, are predicted to cause steric hindrance between FLS2 and BAK1. The Ser residue at position 17 of flg22-ε may cause slight steric hindrance with FLS2.

Together, the motif prevalence, sequence analysis, and protein modelling data led us to hypothesize that most flg22 epitopes of the γ- and β-Proteobacteria are highly immunogenic, whereas most flg22 epitopes of the α-, δ-, and ε-Proteobacteria are weakly to moderately immunogenic. To test this hypothesis, we generated synthetic peptides of the five representative flg22 sequences (**Figure 2**), and monitored their ability to elicit an oxidative burst in *A. thaliana* leaf tissue. Assays using leaf disks from the *bak1-5* mutant (Schwessinger et al. 2011) and accession Ws-4 (a natural *fls2* null mutant) (Gómez-Gómez and Boller 2000) confirmed that the oxidative bursts observed in Col-0 are FLS2/BAK1 dependent (**Figure 4A-E**). Strikingly, and consistent with our hypothesis, only flg22-γ or flg22-β generated oxidative bursts comparable to flg22-Pae in *A. thaliana* Col-0 plants when applied at a concentration of 100 nM (**Figure 4A-B)**. In contrast, the oxidative bursts of *A. thaliana* Col-0 leaves exposed to 100 nM flg22-ε and flg22-δ were approximately 70% and 90% lower, respectively, than those induced by 100 nM flg22-Pae, while no detectable oxidative burst was induced using 100 nM flg22-α (**Figure 4C-E**). Increasing the concentration of flg22-δ led to dose-dependent oxidative bursts, confirming that flg22-δ is weakly immunogenic to *A. thaliana* (**Figure 4F**). On the other hand, no detectable oxidative burst was induced by flg22-α even at 5 µM (**Figure 4F**). However, in seedling growth inhibition assays (**Figure 4G**), 2 µM and 5 µM flg22-α did result in ∼ 20% and ∼ 40% reductions in seedling weight, respectively, suggesting this peptide may weakly associate with the FLS2/BAK1 complex. In support of this, a dose-dependent decrease in 100 nM flg22-Pae-induced oxidative bursts was observed when simultaneously applied to *A. thaliana* Col-0 leaves with 1, 2, or 5 µM of flg22-α as a competitor (**Figure 4H**).

**Figure 4.**
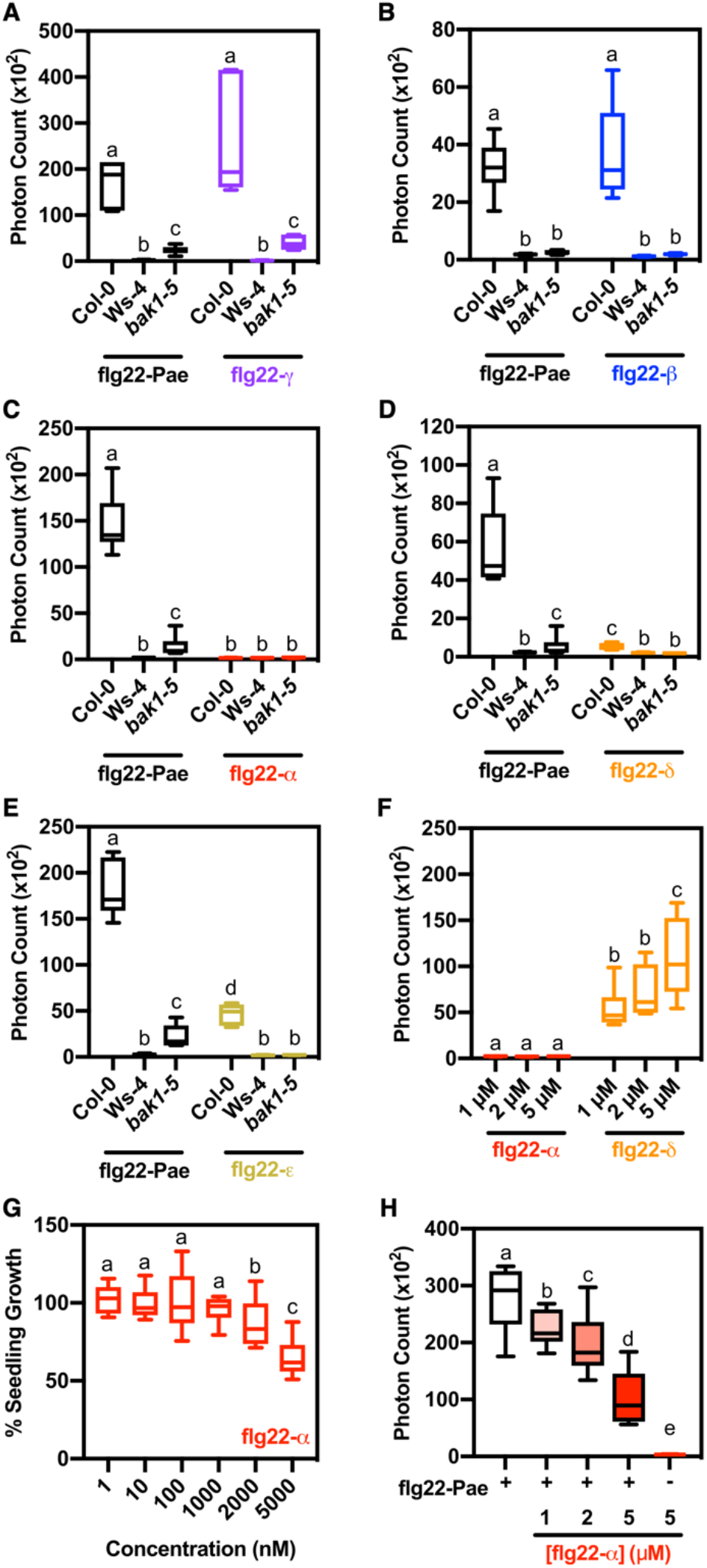
Immunogenicity of proteobacterial flg22 peptides. (**A**-**E**) Oxidative species production in Col-0, *bak1-5* and Ws-4 leaf discs following treatment with 100 nM of the (A) flg22-β, (B) flg22-β, (C) flg22-α, (D) flg22-δ, or (E) flg22-ε peptides. Results for the flg22-Pae peptide are shown in each panel for comparison. Cumulative photon count was measured over 40 mins. (**F**) Flg22-α and flg22-δ-elicited oxidative bursts in Col-0 leaf discs using 1-5 µM of each peptide. (**G**) Seedling growth inhibition of Col-0 seedlings following continual treatment with 1-5000 nM of flg22-α for 14 days. Values are normalized to Col-0 seedlings grown in the absence of flg22. (**H**) Oxidative species production in Col-0 leaf discs following elicitation with 100 nM of flg22-Pae and 1-5 µM of flg22-α. (**A-H**) Data are presented as boxplots indicating first and third quartiles, split by a median line. Whiskers represent maximum and minimum values. Three independent experiments were conducted for all experiments with similar results (n=6).

Our data clearly demonstrate that the immunogenicity of flg22 epitopes varies across the phylum Proteobacteria, with highly immunogenic flg22 epitopes predominantly found in the γ- and β-Proteobacteria. We postulate that this observation relates to the timing of FLS2 evolution relative to the divergence of the proteobacterial classes. The α-Proteobacteria are estimated to have diverged from the γ- and β-Proteobacteria between 2,000 and 3,000 million years ago (mya) (Battistuzzi et al. 2004). In contrast, FLS2 likely emerged following the evolution of land plants (∼ 500 mya) and prior to the diversification of the Spermatophyta (∼ 350 mya) (Albert et al. 2010; de Vries et al. 2018; Morris et al. 2018). We suggest that the divergence of flg22 sequences between proteobacterial classes was not driven by selection to evade plant immune detection, although this may be important in explaining flg22 sequence divergence at finer taxonomic scales. Instead, we hypothesize that the flg22 sequences of the proteobacterial classes were already diverged prior to the emergence of FLS2, and that evolution of FLS2 ligand specificity was driven by the flg22 epitopes of the γ- and β-proteobacteria, a monophyletic group containing many common phytopathogens. However, it is important to recognize that flg22 epitope elicitation varies between plant hosts. In a particularly notable example, the FLS2^XL^ protein of *Vitis riparia* (riverbank grape) triggers a strong immune response following perception of the flg22 epitope of *A. tumefaciens* (Fürst et al. 2020), which is similar in sequence to flg22-α and is a classical example of a “non-immunogenic” flg22 epitope based on studies with *A. thaliana* and tomato (Felix et al. 1999). Similarly, the FLS2 protein of *Glycine max* (soybean) can trigger an immune response upon exposure to the flg22 epitope of *R. solanacearum* (Wei et al. 2020), despite this epitope being non-immunogenic in *A. thaliana* and tomato (Pfund et al. 2004; Mueller et al. 2012). As such, it would be interesting to examine if the five flg22 peptides tested here trigger stronger immune responses in other plant species, and whether there is a relationship between flg22-FLS2 specificity and the host range of the microbes.

## DATA AVAILABILITY

Scripts to repeat the computational analyses reported in this study are available at: github.com/diCenzo-GC/Proteobacterial_flg22.

## ACKNOWLEDGEMENTS

This research was enabled, in part, through computational resources provided by Compute Ontario (computeontario.ca) and Compute Canada (computecanada.ca).

## SUPPLEMENTARY MATERIAL

**File S1**. Supplementary Materials & Methods.

**Figure S1. Distribution of flagellin within the Proteobacteria**. A maximum likelihood phylogeny of 1,414 Proteobacteria is shown, which was prepared from a concatenated alignment of 31 highly conserved proteins. Branches on the tree are colour coded based on taxonomic class, as indicated. Taxa encoding a flagellin predicted to induce a plant immune response, based on the presence of the ^11^[ST]xx[DN][DN]xAGxxI^21^ motif within the flg22, are shown in blue; taxa only encoding flagellin that lack this motif are shown in pink, while taxa lacking flagellin are not coloured. The scale represents the mean number of amino acid substitutions per site.

**Figure S2. Phylogeny of the proteobacterial flagellin proteins**. A maximum likelihood phylogeny of 2,470 flagellin proteins collected from 1,414 Proteobacteria is shown. Ends of the branches on the tree are colour coded based on the taxonomic class of the organism encoding the flagellin, as indicated. The scale represents the mean number of amino acid substitutions per site. Two groups of α-proteobacterial flagellins grouping separately from the rest of the α-proteobacterial flagellins (indicative of horizontal gene transfer) are indicated with arrows and numbers.

**Figure S3. Structural modelling of the FLS2-flg22-BAK1 complex using peptide sequences derived from β- and γ-Proteobacteria**. (**A**) Association between the amino-terminal portion of flg22 (positions 1-9) (red) and FLS2 (purple). The flg22 epitope encoded by *P. aeruginosa* (Pae) and FLS2 is stabilized by H-bonds (dotted yellow lines) between the backbone of flg22 at Gln-1 and Leu-3 and the side chains of Tyr-148 and Arg-152 of FLS2, respectively (Sun et al. 2013). H-bonding is predicted at these positions for all modelled peptides. (**B**) Interaction between the central portion of flg22 (positions 10-15) and FLS2 is stabilized by H-bonds between Tyr-272, Arg-294, Tyr-296, and His-316 of FLS2 and Leu-13 and Asp-14 of flg22 (Sun et al. 2013), which is predicted for all flg22 peptides. Asp-14 and Asp-15 of flg22-Pae interact with two positively charged pockets on the surface of FLS2 (blue) (Sun et al. 2013). (**C**) The carboxyl-terminal portion of flg22 (positions 16-22) interacts with FLS2 and BAK1 (green). Leu-19 of flg22 forms H-bonds with Thr-52 and Val-54 of BAK1 (Sun et al. 2013) and interacts with a hydrophobic pocket (red) on the surface of FLS2 (Sun et al. 2013).

**Table S1**. Distribution of flagellins containing the ^11^[ST]xx[DN][DN]xAGxxI^21^ motif within flg22.

**Table S2**. Predicted changes in flg22 binding affinity to FLS2, and of flg22-bound FLS2 to BAK1.

**Dataset S1. Metadata of the Proteobacteria analyzed in this study**. The metadata (as provided by the NCBI Genome database) for the 1,414 proteobacterial strains used in this study are provided. All species are named according to their name in the NCBI Genome database at the time of download, which may not necessarily be the currently accepted name. Data for each class is provided as a separate worksheet, with the exception of the δ-Proteobacteria and the ε-Proteobacteria, which are combined as these classes are combined in the NCBI Genome database.

## REFERENCES

Albert, M. K. Jehle, A., Lipschis, M., Mueller, K., Zeng, Y., and Felix, G. 2010. Regulation of cell behaviour by plant receptor kinases: Pattern recognition receptors as prototypical models. Eur. J. Cell Biol. 89:200–207

Ausubel, F. M. 2005. Are innate immune signaling pathways in plants and animals conserved? Nat. Immunol. 6:973–979

Battistuzzi, F. U., Feijao, A., and Hedges, S. B. 2004. A genomic timescale of prokaryote evolution: insights into the origin of methanogenesis, phototrophy, and the colonization of land. BMC Evol. Biol. 4:44

Bauer, Z., Gómez-Gómez, L., Boller, T., and Felix, G. 2001. Sensitivity of different ecotypes and mutants of Arabidopsis thaliana toward the bacterial elicitor flagellin correlates with the presence of receptor-binding sites. J. Biol. Chem. 276:45669–45676

Chinchilla, D., Zipfel, C., Robatzek, S., Kemmerling, B., Nürnberger, T., Jones, J. D. G., Felix, G., and Boller, T. 2007. A flagellin-induced complex of the receptor FLS2 and BAK1 initiates plant defence. Nature. 448:497–500

Couto, D., and Zipfel, C. 2016. Regulation of pattern recognition receptor signalling in plants. Nat. Rev. Immunol. 16:537–552

de Vries, S., de Vries, J., von Dahlen, J. K., Gould, S. B., Archibald, J. M., Rose, L. E., and Slamovits, C. H. 2018. On plant defense signaling networks and early land plant evolution. Commun. Integr. Biol. 11:1–14

Felix, G., Duran, J. D., Volko, S., and Boller, T. 1999. Plants have a sensitive perception system for the most conserved domain of bacterial flagellin. Plant J. 18:265–276

Fürst, U., Zeng, Y., Albert, M., Witte, A. K., Fliegmann, J., and Felix, G. 2020. Perception of Agrobacterium tumefaciens flagellin by FLS2^XL^ confers resistance to crown gall disease. Nat. Plants. 6:22–27

Gómez-Gómez, L., and Boller, T. 2000. FLS2: an LRR receptor–like kinase involved in the perception of the bacterial elicitor flagellin in Arabidopsis. Mol. Cell. 5:1003–1011

Gourion, B., Berrabah, F., Ratet, P., and Stacey, G. 2015. Rhizobium-legume symbioses: the crucial role of plant immunity. Trends Plant Sci. 20:186–194

Gust, A. A., Pruitt, R., and Nürnberger, T. 2017. Sensing danger: key to activating plant immunity. Trends Plant Sci. 22:779–791

Morris, J. L., Puttick, M. N., Clark, J. W., Edwards, D., Kenrick, P., Pressel, S., Wellman, C. H., Yang, Z., Schneider, H., and Donoghue, P. C. J. 2018. The timescale of early land plant evolution. Proc. Natl. Acad. Sci. U.S.A. 115:E2274–E2283

Mueller, K., Bittel, P., Chinchilla, D., Jehle, A. K., Albert, M., Boller, T., and Felix, G. 2012. Chimeric FLS2 receptors reveal the basis for differential flagellin perception in Arabidopsis and tomato. Plant Cell. 24:2213–2224

Nicaise, V., Roux, M., and Zipfel, C. 2009. Recent advances in PAMP-triggered immunity against bacteria: pattern recognition receptors watch over and raise the alarm. Plant Physiol. 150:1638–1647

Pfund, C., Tans-Kersten, J., Dunning, F. M., Alonso, J. M., Ecker, J. R., Allen, C., and Bent, A. F. 2004. Flagellin is not a major defense elicitor in Ralstonia solanacearum cells or extracts applied to Arabidopsis thaliana. Mol. Plant Microbe Interact. 17:696–706

Plett, J. M., and Martin, F. M. 2018. Know your enemy, embrace your friend: using omics to understand how plants respond differently to pathogenic and mutualistic microorganisms. Plant J. 93:729–746

Schwessinger, B., Roux, M., Kadota, Y., Ntoukakis, V., Sklenar, J., Jones, A., and Zipfel, C. 2011. Phosphorylation-dependent differential regulation of plant growth, cell death, and innate immunity by the regulatory receptor-like kinase BAK1. PLOS Genet. 7:e1002046

Sun, W., Dunning, F. M., Pfund, C., Weingarten, R., and Bent, A. F. 2006. Within-species flagellin polymorphism in Xanthomonas campestris pv campestris and its impact on elicitation of Arabidopsis FLAGELLIN SENSING2–dependent defenses. Plant Cell. 18:764–779

Sun, Y., Li, L., Macho, A. P., Han, Z., Hu, Z., Zipfel, C., Zhou, J.-M., and Chai, J. 2013. Structural basis for flg22-induced activation of the Arabidopsis FLS2-BAK1 immune complex. Science. 342:624–628

Trdá, L., Fernandez, O., Boutrot, F., Héloir, M.-C., Kelloniemi, J., Daire, X., Adrian, M., Clément, C., Zipfel, C., Dorey, S., and Poinssot, B. 2014. The grapevine flagellin receptor VvFLS2 differentially recognizes flagellin-derived epitopes from the endophytic growth-promoting bacterium Burkholderia phytofirmans and plant pathogenic bacteria. New Phytol. 201:1371–1384

Wei, Y., Balaceanu, A., Rufian, J. S., Segonzac, C., Zhao, A., Morcillo, R. J. L., and Macho, A. P. 2020. An immune receptor complex evolved in soybean to perceive a polymorphic bacterial flagellin. Nat. Commun. 11:3763

Wheeler, T. J., Clements, J., and Finn, R. D. 2014. Skylign: a tool for creating informative, interactive logos representing sequence alignments and profile hidden Markov models. BMC Bioinform. 15:7

Zhang, N., Chen, Y., Lu, H., Zhao, F., Alvarez, R. V., Goncearenco, A., Panchenko, A. R., and Li, M. 2020. MutaBind2: predicting the impacts of single and multiple mutations on protein-protein interactions. iScience. 23:100939

