## Supplemental File S1 for "Proteobacteria encode diverse flg22 peptides that elicit varying immune responses in *Arabidopsis thaliana*"

### SUPPLEMENTARY MATERIAL AND METHODS

#### Plant growth conditions

*Arabidopsis thaliana* ecotype Columbia-0 (Col-0) and Wassilewskjia-4 (Ws-4) were sown on moist soil and allowed to germinate. One week after germination, six seedlings of each genotype were carefully transplanted to 25 cm x 15 cm pots containing Sunshine Mix #1 (SunGro) soil and allowed to grow for four additional weeks under short-day conditions (8 h light; 150 μE $m^{-2}$ $s^{-1}$) at 22°C in a climate-controlled growth chamber (BioChambers). Soil was supplemented with 1.5 g/L standard 20:20:20 N:P:K fertilizer two weeks after transplanting to larger pots, and predatory *Amblyseius swirskii* mites (Koppert Biological Systems) were released into the chambers twice during the growth chamber as a precaution against glasshouse pests.

#### Immune assays

Peptides used in immune assays were synthesized by EZBiolab (Parsippany, New Jersey, USA) with 95% purity, and resuspended to a concentration of 10 mM in pure water. Oxidative burst and seedling growth inhibition assays were conducted as described in Bredow et al. (2019).

#### Genomic dataset

The RefSeq version of 1,414 proteobacterial proteomes and Genbank flat files (366 α-Proteobacteria, 234 β-Proteobacteria, 674 γ-Proteobacteria, 74 δ-Proteobacteria, and 66 ε-Proteobacteria) were downloaded from the National Center for Biotechnology Information (NCBI) Genome database between 17 June 2020 and 31 July 2020. Strains were included in our analysis if they had a genome assembly level of 'Complete' or 'Chromosome', and if they belonged to the 'Reference' or 'Representative' RefSeq categories. A complete list of strains, and associated metadata, is provided as Dataset S1.

#### Identification and sequence analysis of flagellin proteins

Flagellin proteins were extracted from the 1,414 Proteobacteria using an adaptation of our previously described pipeline (diCenzo et al. 2019). The analysis was run separately for each proteobacterial class, with the exception that the Deltaproteobacteria and Epsilonproteobacteria were analyzed together as these datasets are combined in NCBI. Briefly, the Flagellin_N hidden Markov model (HMM) of the Pfam database (El-Gebali et al. 2019) was used to quickly identify putative flagellin proteins using the HMMSEARCH function of HMMER 3.3 (Eddy 2009). All hits, regardless of the score, were collected and searched against a combined HMM database containing 18,259 HMMs from Pfam 33.1, 4,490 HMMs from TIGRFAM 15.0 (Haft et al. 2013), and HMMs of the PRK06008 and PRK12687 conserved protein domain families (Lu et al. 2020) using the HMMSCAN function of HMMER. The top scoring HMM hit for each query protein was identified, and proteins whose top scoring hit was either Flagellin_N, Flagellin_C, or PRK12687 were annotated as flagellin.

All flagellin within a Proteobacterium class were aligned using Clustal Omega 1.2.4 (Sievers et al. 2011), and the alignments manually scanned to identify the flg22 epitopes. The flg22 epitope of each protein was extracted, and all sequences containing indels were discarded. Amino acid logos of the flg22 regions were prepared using Skylign with the 'use observed counts' and 'information content – above background' options (Wheeler et al. 2014). Flg22 epitopes containing a [ST]xx[DN][DN]xAGxxI motif were identified using Grep, and subsequently confirmed to occur between position 11 and 21 in all of the flg22 epitopes containing this motif.

To examine their evolutionary relationships, the 2,470 flagellin proteins identified across the 1,414 Proteobacteria were combined as a single dataset, aligned using Clustal Omega 1.2.4 (Sievers et al. 2011), and trimmed using TRIMAL 1.4rev15 with the automated1 option (Capella-Gutiérrez et al. 2009). The alignment was used to construct a maximum likelihood phylogeny using raxmlHPC-HYBRID-AVC 8.2.12 (Stamatakis 2014) with the CAT model of rate heterogeneity and the LG amino acid substitution model, with bootstrapping stopped based on the extended majority-rule consensus tree criterion. The LG amino acid substitution model was chosen as it gave the best scoring trees in a preliminary test of model options. Phylogenies were prepared on a Compute Canada server, and they were visualized with the iTOL web server (Letunic and Bork 2016).

**Protein structure modelling and binding affinity predictions**

The crystal structure of FLS2-flg22-BAK1 (PDB:4MN8) (Sun et al. 2013) was modelled using PyMOL Molecular Graphics System Version 2.4.0. Flg22 peptides representative of each proteobacterial class were generated using the PyMOL Mutagenesis Wizard. Surface charges were predicted using the APBS Electrostatics Plugin (prepwizard: SCHRODINGER method). The prediction of hydrogen bonds between flg22 and FLS2/BAK1 was elucidated using the find polar contacts function.

Total changes in binding affinity between flg22 peptides and FLS2/BAK1 were calculated using MutaBind2 (Zhang et al. 2020) and the crystal structure of the FLS2-flg22-BAK1 complex (PDB:4MN8) (Sun et al. 2013). Binding affinities were estimated separately for phase 1 (FLS2-flg22) and phase 2 (FLS2-flg22-BAK1) of complex formation.

**Species phylogenetic analyses**

An unrooted phylogeny of the 1,414 Proteobacteria was constructed using an adaptation of our previously described pipeline (diCenzo et al. 2019). Briefly, a set of 31 single-copy proteins (DnaG, Frr, InfC, NusA, Pgk, PyrG, RplA, RplB, RplC, RplD, RplE, RplF, RplK, RplL, RplM, RplN, RplP, RplS, RplT, RpmA, RpoB, RpsB, RpsC, RpsE, RpsI, RpsJ, RpsK, RpsM, RpsS, SmpB, Tsf) found in at least 95% of the proteomes were identified using the AMPHORA2 pipeline (Wu and Scott 2012) and custom scripts. Orthologous groups were aligned using MAFFT 7.467 with the localpair option (Katoh and Standley 2013), following which the alignments were trimmed using TRIMAL 1.4rev15 with the automated1 option (Capella-Gutiérrez et al. 2009). Alignments were concatenated and used to construct a maximum likelihood phylogeny using raxmlHPC-HYBRID-AVC 8.2.12 (Stamatakis 2014) with the CAT model of rate heterogeneity and the LG amino acid substitution model, with 100 bootstrap replicates. The LG amino acid substitution model was chosen as it gave the best scoring trees in a preliminary test of model options. Phylogenies were prepared on a Compute Canada server, and they were visualized with the iTOL web server (Letunic and Bork 2016).

**Data availability**

Scripts to repeat the computational analyses reported in this study are available at github.com/diCenzo-GC/Proteobacterial_flg22.

### SUPPLEMENTARY REFERENCES


Bredow, M., Sementchoukova, I., Siegel, K., and Monaghan, J. 2019. Pattern-triggered oxidative burst and seedling growth inhibition assays in *Arabidopsis thaliana*. J. Vis. Exp. :e59437

Capella-Gutiérrez, S., Silla-Martínez, J. M., and Gabaldón, T. 2009. trimAl: a tool for automated alignment trimming in large-scale phylogenetic analyses. Bioinformatics. 25:1972–1973

diCenzo, G. C., Mengoni, A., and Perrin, E. 2019. Chromids aid genome expansion and functional diversification in the family *Burkholderiaceae*. Mol. Biol. Evol. 36:562–574

Eddy, S. R. 2009. A new generation of homology search tools based on probabilistic inference. Genome Inform. 23:205–211

El-Gebali, S., Mistry, J., Bateman, A., Eddy, S. R., Luciani, A., Potter, S. C., Qureshi, M., Richardson, L. J., Salazar, G. A., Smart, A., Sonnhammer, E. L. L., Hirsh, L., Paladin, L., Piovesan, D., Tosatto, S. C. E., and Finn, R. D. 2019. The Pfam protein families database in 2019. Nucleic Acids Res. 47:D427–D432

Haft, D. H., Selengut, J. D., Richter, R. A., Harkins, D., Basu, M. K., and Beck, E. 2013. TIGRFAMs and genome properties in 2013. Nucleic Acids Res. 41:D387–D395

Katoh, K., and Standley, D. M. 2013. MAFFT multiple sequence alignment software version 7: improvements in performance and usability. Mol. Biol. Evol. 30:772–780

Letunic, I., and Bork, P. 2016. Interactive tree of life (iTOL) v3: an online tool for the display and annotation of phylogenetic and other trees. Nucleic Acids Res. 44:W242–W245

Lu, S., Wang, J., Chitsaz, F., Derbyshire, M. K., Geer, R. C., Gonzales, N. R., Gwadz, M., Hurwitz, D. I., Marchler, G. H., Song, J. S., Thanki, N., Yamashita, R. A., Yang, M., Zhang, D., Zheng, C., Lanczycki, C. J., and Marchler-Bauer, A. 2020. CDD/SPARCLE: the conserved domain database in 2020. Nucleic Acids Res. 48:D265–D268

Sievers, F., Wilm, A., Dineen, D., Gibson, T. J., Karplus, K., Li, W., Lopez, R., McWilliam, H., Remmert, M., Söding, J., Thompson, J. D., and Higgins, D. G. 2011. Fast, scalable generation of high-quality protein multiple sequence alignments using Clustal Omega. Mol. Syst. Biol. 7:539

Stamatakis, A. 2014. RAxML version 8: a tool for phylogenetic analysis and post-analysis of large phylogenies. Bioinformatics. 30:1312–1313

Sun, Y., Li, L., Macho, A. P., Han, Z., Hu, Z., Zipfel, C., Zhou, J.-M., and Chai, J. 2013. Structural basis for flg22-induced activation of the *Arabidopsis* FLS2-BAK1 immune complex. Science. 342:624–628

Wheeler, T. J., Clements, J., and Finn, R. D. 2014. Skylign: a tool for creating informative, interactive logos representing sequence alignments and profile hidden Markov models. BMC Bioinform. 15:7

Wu, M., and Scott, A. J. 2012. Phylogenomic analysis of bacterial and archaeal sequences with AMPHORA2. Bioinformatics. 28:1033–1034

Zhang, N., Chen, Y., Lu, H., Zhao, F., Alvarez, R. V., Goncearenco, A., Panchenko, A. R., and Li, M. 2020. MutaBind2: predicting the impacts of single and multiple mutations on protein-protein interactions. iScience. 23:100939
