## Supplemental Figure S1 for "Proteobacteria encode diverse flg22 peptides that elicit varying immune responses in *Arabidopsis thaliana*"

**Figure S1. Distribution of flagellin within the Proteobacteria.** A maximum likelihood phylogeny of 1,414 Proteobacteria is shown, which was prepared from a concatenated alignment of 31 highly conserved proteins. Branches on the tree are colour coded based on taxonomic class, as indicated. Taxa encoding a flagellin predicted to induce a plant immune response, based on the presence of the $_{11}$[ST]xx[DN][DN]xAGxxI$_{21}$ motif within the flg22, are shown in blue; taxa only encoding flagellin that lack this motif are shown in pink, while taxa lacking flagellin are not coloured. The scale represents the mean number of amino acid substitutions per site.

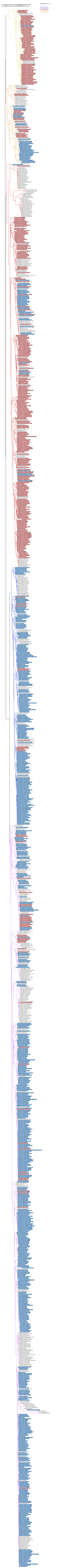
