## Supplemental Figure S2 for "Proteobacteria encode diverse flg22 peptides that elicit varying immune responses in *Arabidopsis thaliana*"

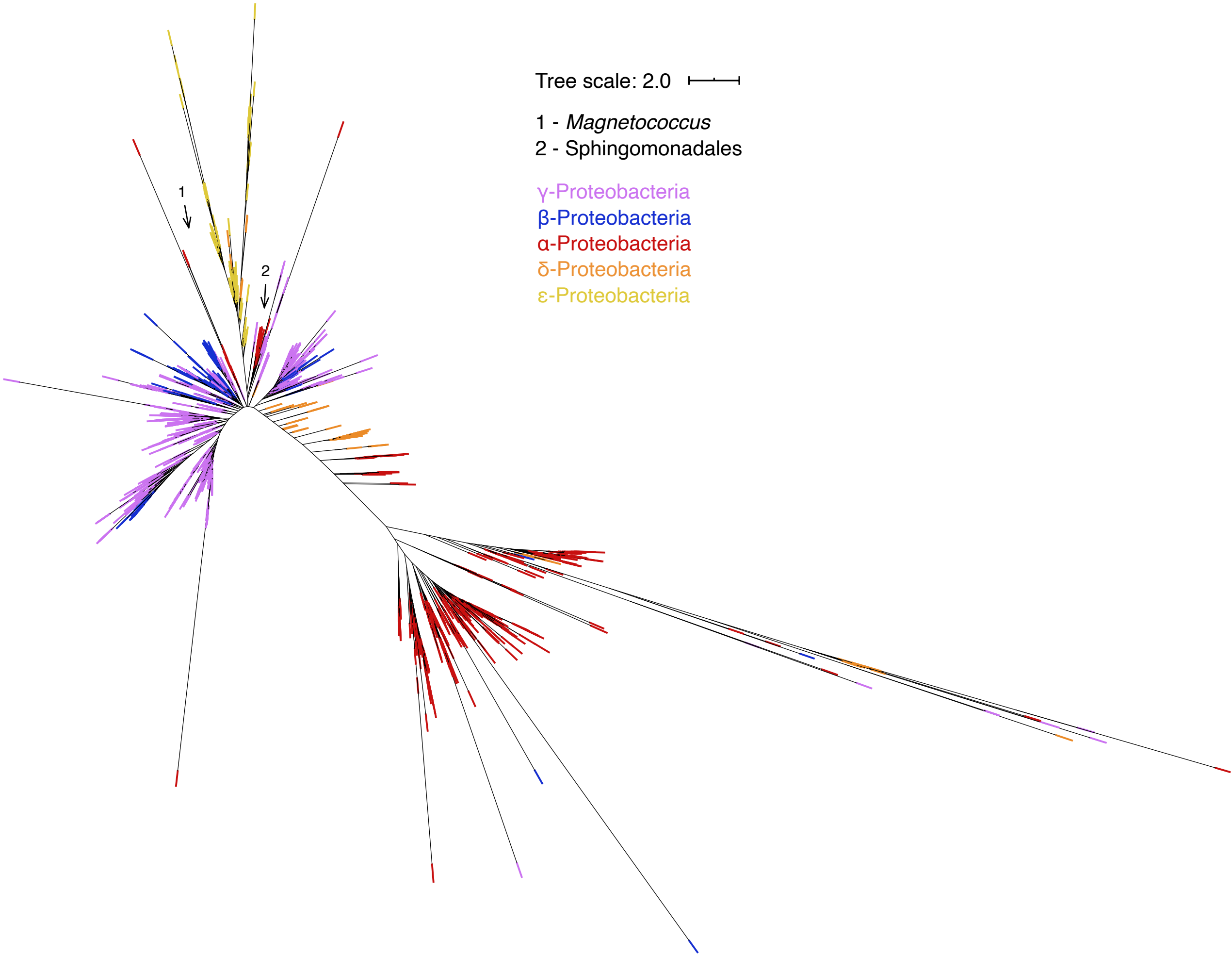


**Figure S2. Phylogeny of the proteobacterial flagellin proteins.** A maximum likelihood phylogeny of 2,470 flagellin proteins collected from 1,414 Proteobacteria is shown. Ends of the branches on the tree are colour coded based on the taxonomic class of the organism encoding the flagellin, as indicated. The scale represents the mean number of amino acid substitutions per site. Two groups of α-proteobacterial flagellins grouping separately from the rest of the α-proteobacterial flagellins (indicative of horizontal gene transfer) are indicated with arrows and numbers.
