## Supplemental Figure S3 for "Proteobacteria encode diverse flg22 peptides that elicit varying immune responses in *Arabidopsis thaliana*"

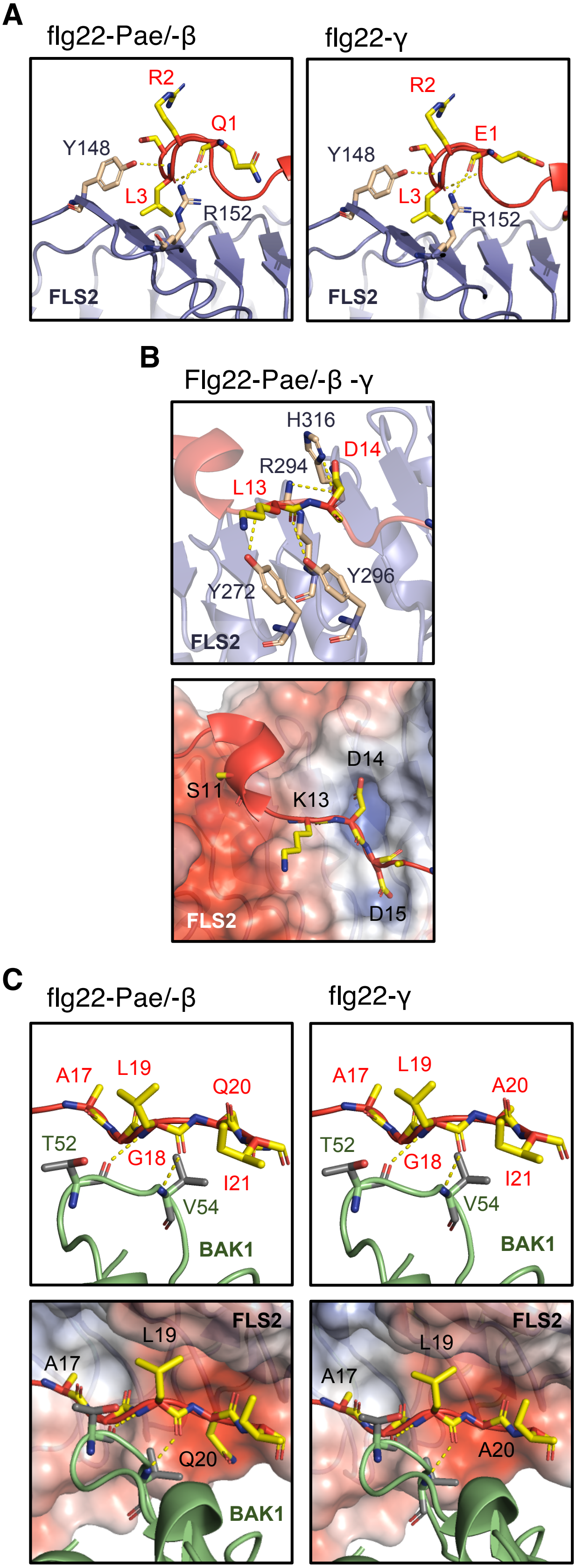


**Figure S3. Structural modelling of the FLS2-flg22-BAK1 complex using peptide sequences derived from β- and γ-Proteobacteria.** (**A**) Association between the amino-terminal portion of flg22 (positions 1-9) (red) and FLS2 (purple). The flg22 epitope encoded by P. aeruginosa (Pae) and FLS2 is stabilized by H-bonds (dotted yellow lines) between the backbone of flg22 at Gln-1 and Leu-3 and the side chains of Tyr-148 and Arg-152 of FLS2, respectively (Sun et al. 2013). H-bonding is predicted at these positions for all modelled peptides. (**B**) Interaction between the central portion of flg22 (positions 10-15) and FLS2 is stabilized by H-bonds between Tyr-272, Arg-294, Tyr-296, and His-316 of FLS2 and Leu-13 and Asp-14 of flg22 (Sun et al. 2013), which is predicted for all flg22 peptides. Asp-14 and Asp-15 of flg22-Pae interact with two positively charged pockets on the surface of FLS2 (blue) (Sun et al. 2013). (**C**) The carboxyl-terminal portion of flg22 (positions 16-22) interacts with FLS2 and BAK1 (green). Leu-19 of flg22 forms H-bonds with Thr-52 and Val-54 of BAK1 (Sun et al. 2013) and interacts with a hydrophobic pocket (red) on the surface of FLS2 (Sun et al. 2013).
