## Supplemental Table S1 for "Proteobacteria encode diverse flg22 peptides that elicit varying immune responses in *Arabidopsis thaliana*"

**Table S1.** Distribution of flagellins containing the [11][ST]xx[DN][DN]xAGxxI[21] motif within flg22.

| | α-Proteobacteria | β-Proteobacteria | γ-Proteobacteria | δ-Proteobacteria | ε-Proteobacteria |
|---|---|---|---|---|---|
| Total strains * | 366 | 234 | 675 | 74 | 66 |
| Strains with flagellin † | 264 (72%) | 188 (80%) | 508 (75%) | 49 (66%) | 61 (92%) |
| Strains with motif ¥ | 39 (15%) | 178 (95% | 469 (92%) | 29 (59%) | 1 (2%) |

* The total number of strains of each proteobacterial class included in the analysis.

† The number of strains of each proteobacterial class encoding at least one flagellin. Values in the parenthesis indicate the percentage of all strains that encode at least one flagellin.

¥ The number of strains of each proteobacterial class encoding at least one flagellin containing the [11][ST]xx[DN][DN]xAGxxI[21] motif in the flg22 domain. Values in the parenthesis indicate the percentage of strains encoding at least one flagellin that encode a flagellin with this motif.
