## Supplemental Table S2 for "Proteobacteria encode diverse flg22 peptides that elicit varying immune responses in *Arabidopsis thaliana*"

**Table S2.** Flg22 peptide binding affinity estimates to FLS2 or BAK1 calculated using the MutBind2 server.

| Peptide | Sequence | Phase 1 (FLS2-flg22) | Phase 2 (FLS2/flg22-BAK1) |
|---|---|---|---|
| | | $\Delta\Delta G_{bind}$ (kcal/mol) | $\Delta\Delta G_{bind}$ (kcal/mol) |
| flg22-γ | ERLSSGLRINSAKDDAAGLQIS | -0.91 | -0.91 |
| flg22-β | QRLSSGLRINSAKDDAAGLAIA | -0.81 | -0.54 |
| flg22-α | NRISTGKRVNSASDNAAYWSIA | 2.37 | 1.37 |
| flg22-δ | RRLSSGLRINTAADDAAGLAIR | 0.32 | 0.29 |
| flg22-ε | EKLSSGLRINKAADDASGLAIA | 0.81 | 0.18 |
